# Understanding distribution and occupancy of Himalayan Monal in Uttarkashi district, Uttarakhand

**DOI:** 10.1101/2021.02.16.431367

**Authors:** Amira Sharief, Hemant Singh, Bheem Dutt Joshi, Tanoy Mukherjee, Kailash Chandra, Mukesh Thakur, Lalit Kumar Sharma

**Affiliations:** Zoological Survey of India, New Alipore, Kolkata, 700053 West Bengal, India; Wildlife Institute of India, Dehradun, 248001 Uttarakhand, India

**Keywords:** Himalayan monal, occupancy, abundance, activity pattern, Western Himalayas

## Abstract

The Himalayan Monal is a conservation priority species in its entire distribution range. Its population is declining in many areas due to various anthropogenic threats. The information on species distribution and its abundance is lacking in many areas which are vital for conservation and management planning. Hence, through the present study, we have assessed the abundance and occupancy of Himalayan monal in Uttarkashi district (Uttarakhand). We used camera traps and conventional sign surveys for documenting the species during 2018-2019. We installed a total of 69 camera traps (2819 trap nights) and surveyed 54 trails (650 km) which represents entire habitat and topographic variability of the landscape. The occupancy and detection probability was modelled using the habitat variables. The top model showed that occupancy probability of Himalayan monal was positively influenced by the slope (β =27.52 ±16.25) and negatively influenced by Reserve Forest (RF) (β= −8.14 SE ± 4.99). The observed naïve occupancy of Himalayan Monal was 0.69 in the study area, which was slightly lower than the estimated occupancy. However, in the null model, the site occupancy estimated was found to be 0.82±0.08 and with detection probability 0.23±0.03. The overall abundance of monal was estimated about 171.58 ±10.2 in the study area with an average density of 0.62/ km^2^. The activity pattern analysis indicates that monal remains very active between 6.00 hrs −12.00 hrs and relatively less active during mid-day when humans are most active 11.30 hrs-16.30 hrs. The present study is a first attempt to estimate occupancy and abundance using camera traps as well as sign survey for the species primarily from non-Protected Area (PA). We found that Himalayan monal is abundant outside the PAs, which is a good indication for its long-term viability and also identified areas for conservation and management prioritization in Uttarkashi.

## Introduction

The Himalayan bio-geographic zone is situated at the junction of three bio-geographical realms viz., Palaearctic, Africo-tropical and Indo-Malayan and home to some of the top conservation priority fauna Mani (1974). Its rich habitat and climatic variability have resulted in the creation of safe adobe for over 30,000 different faunal species Chandra et al (2018). The Pheasants belong to order Galliformes, a group of charismatic animals among all the faunal elements distributed in the Himalayan landscape. These pheasant species found in India are most abundant in the middle and high altitude valleys of the Himalayan range, and the majority of them are endemic to Himalayas Sathyakumar et al (2011). They are very sensitive to anthropogenic disturbance and habitat degradation Fuller & Garson (2000), hence also considered as bio-indicators, they also form a prey base of carnivores birds and mammals Johnsgard (1986). These birds are popular in folklore and have been used for conservation campaigns as icons because of their importance in ecosystem and unique breeding displays Nawaz et al (2000). Having a distinct and unique colouration pattern, pheasant species can easily be differentiated from other species of birds Ali (1981). Of the 51 species of 16 genera of pheasants found in Asia, of which 17 species are reported to be present in India Ali & Ripley (1987). Out of 17 species, 16 species of pheasants are reported from the Himalaya, among which Himalaya beholds five threatened species Sathyakumar & Kaul (2007). These birds are reported from diverse habitats from lowlands tropical forests to temperate coniferous forest.

Most of the pheasant species are expanding their ranges to subalpine scrub, alpine meadows, montane grass scrub, and broad-leaved evergreen forests up to the highest physiologically possible elevation in the Himalayas. The Himalayan monal (*Lophophorus impejanus*) a bird with beautiful plumage and with prominent sexual dimorphism is distributed in the upper-middle elevations of Himalayas Delacour (1977). Hence, because of the species is iconic and is off conservation importance, these birds have been given the status of National bird of Nepal and State bird of Uttarakhand state of India. It occupies the montane ecosystem of Himalaya from eastern Afghanistan, Pakistan, India, Nepal, Bhutan, China and Myanmar Sathyakumar & Kaul (2007); BirdLife-International (2020). The species mostly occupies the upper temperate forests of conifer and oak with open grasslands slopes between the elevational ranges of 2400 – 4500 m Grimmett et al (1998). Researchers have observed that Himalayan monal exhibited with seasonal migration along with the altitudinal gradient Gaston & Garson (1981), mostly distributed at altitudes between 2620 m and 3350 m in summer and between 2000 m and 2800 m in winter, with relative preference to the sub-alpine oak forest in spring and conifer, dominated forest during winter Ramesh (2003).

In the entire Himalayan range, recent developmental activities, high level of human disturbance, increase in livestock population has led to degradation and deforestation of forested habitats Bhattacharya et al (2009); Jolli & M. Pandit (2011). The reductions in the forested area, along with the fragmentation of habitat, has adversely impacted the pheasant’s distribution and population abundance Diamond (1974); Sathyakumar (2007); Ramesh (2003); Singh et al (2011). Habitat loss and degradation, hunting for food, sport & trade and human disturbance as other major threats of Himalayan monal and other pheasants throughout their distribution range Fuller & Garson (2000). Despite this, limited information is available on the pheasant population biology and behaviour due to their elusive nature, and rugged terrain in their habitat makes them difficult to observe. Hence, conventional methods did not yield significant information necessary for making strategies for conservation and management. However, camera traps have been found useful in documenting these birds distribution and abundance Dinata et al (2008); Li et al (2010); Suwanrat et al (2015). Nevertheless, more information is needed on the species distribution, abundance and other ecological aspects of the species for making informed management strategies throughout its entire range.

The information available on this species is mostly scattered and focused on Protected Areas (PA) that is considered the best habitats of the species. However, no information is available from forested habitats outside the PA network, which may be acting as a biological corridor connecting PAs. There are large size PAs *viz*., Gangotri National Park (GNP), Govind Pashu Vihar National Park (GPVNP) and Govind Pashu Vihar Wildlife Sanctuary (GPVWLS) are located in Uttarkashi district which possess Himalayan monal populations. Still, the non-PA or the territorial forest/reserve forest between these PAs have never been assessed. However, this area may be acting as a biological corridor or mediating gene flow which is vital for the viability of Himalayan monal population in PAs. Therefore, we designed the present study aiming at assessing the distribution, site occupancy, abundance estimation and activity pattern analysis of Himalayan monal in forested habitats outside PA network in Uttarkashi district of Uttarakhand. We hypothesised that human disturbance will impact the Himalayan monal occupancy and its activity pattern Nawaz et al (2000). Further we attempted to understand the key habitat predictors and their influence on occupancy of Himalayan monal in the study landscape.

### Study area

The study was conducted in the Uttarkashi district of Uttarakhand (Fig. 1), which is located between 38° 28’ to 31°28’ N latitude and 77°49’ to 79°25’ E longitude. The district, covering an area of 8016 km^2^, is located in the north-western region of Indian Himalayan Region at an elevation of 1158–6323 m. The climate of the district varies according to the altitude and aspect and is temperate monsoonal type. The terrain is mountainous consisting of hill ranges and narrow valleys. The district lies in the upper catchment of two major rivers of India *viz*., Ganges (Bhagirathi towards upstream) and the Yamuna. The dominant vegetation types of the study area are Himalayan moist temperate forest, sub-alpine forest and alpine scrub (Champion & Seth 1968). Broad land use categories of the area include permanent settlements (villages), irrigated and rainfed agricultural fields, scrubland, mixed broadleaf forests, subalpine oak-fir forest, summer camping sites and alpine meadows locally known as “Kharaks” and “bugyals” (Awasthi et al. 2003). The entire district is divided into three Forest Divisions; i) Uttarkashi Forest Division ii) Upper Yamuna Forest Division iii) Tons Forest Division and also has three PAs *viz*., GNP, GPVNP and GPVWLS. The forested habitats of the study landscape are home to top conservation priority species including Snow Leopard, Musk deer, Asiatic black bear, Himalayan Brown bear Western Tragopan and Himalayan monal.

**Fig. 1.**
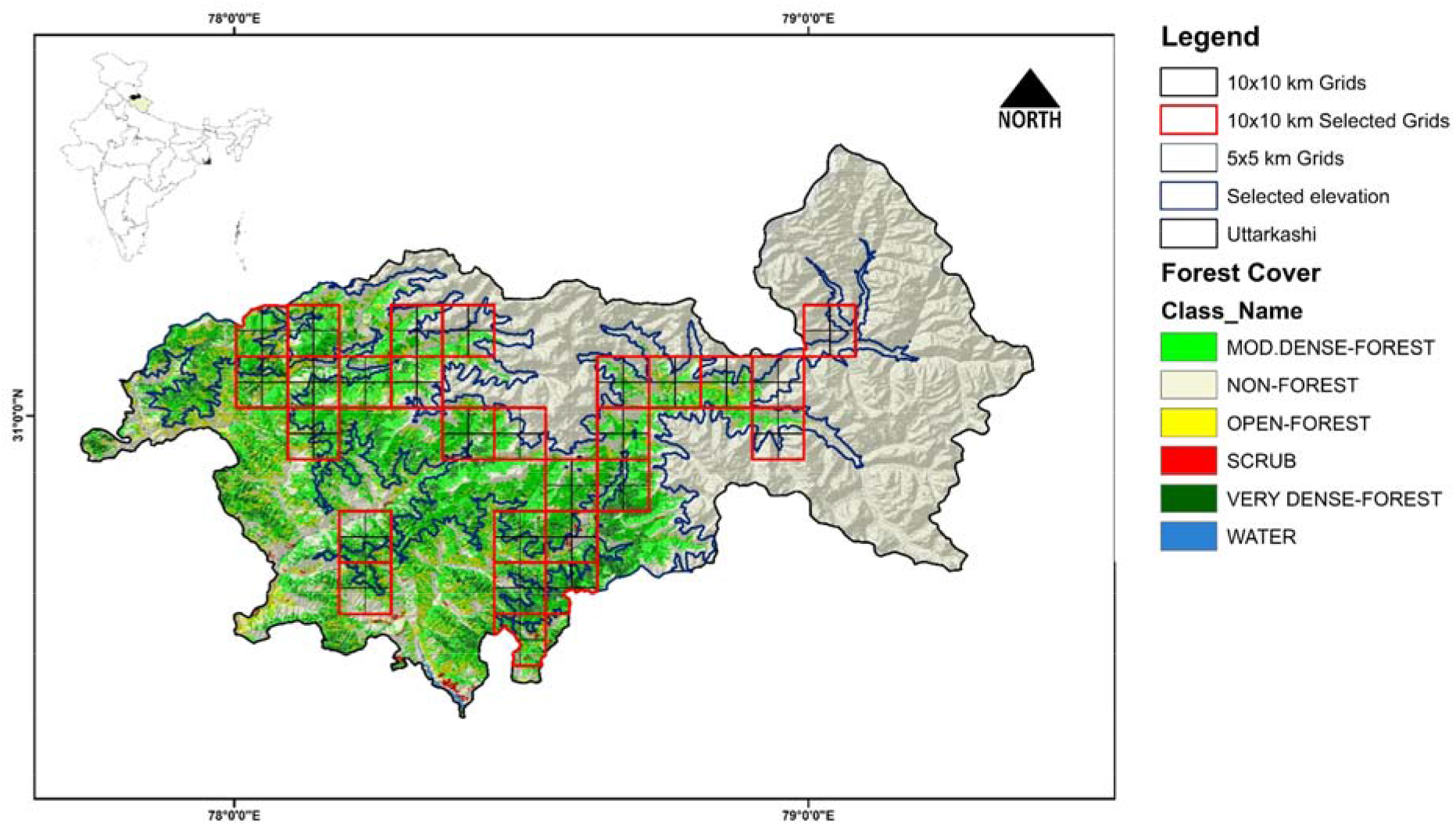
Map of study area Uttarkashi, Uttarakhand.

## Methodology

### Data collection

For collecting information on Himalayan monal, we first stratified the study landscape into different forest types and cover (Fig. 1 and Table 1). The forest types were identified based on the forest types classified by the Forest Survey of India (ISFR 2019). The entire study area was divided into 10 km × 10 km grids to maximize our effort with an aim to cover all logistically accessible grids. The field surveys were conducted during 2018-2019 in all the different habitats (dense forest, open forest, alpine meadows, near human habitations, community and reserved forest) of species in the species distribution elevation range of Uttarkashi district. We used a two-pronged approach, *i*.*e*. transect surveys and camera trapping, and we conducted our study in the 26 selected grids of 10 km × 10 km size which possess species habitat out of the total 133 grids in the elevation gradient (2000–4200 m; Fig. 1). A reconnaissance survey was conducted in all the selected grids for documenting the species presence. All direct or indirect signs (direct sighting, shed feature) of Himalayan monal were recorded during the surveys by the team of three researchers.

**Table 1:**
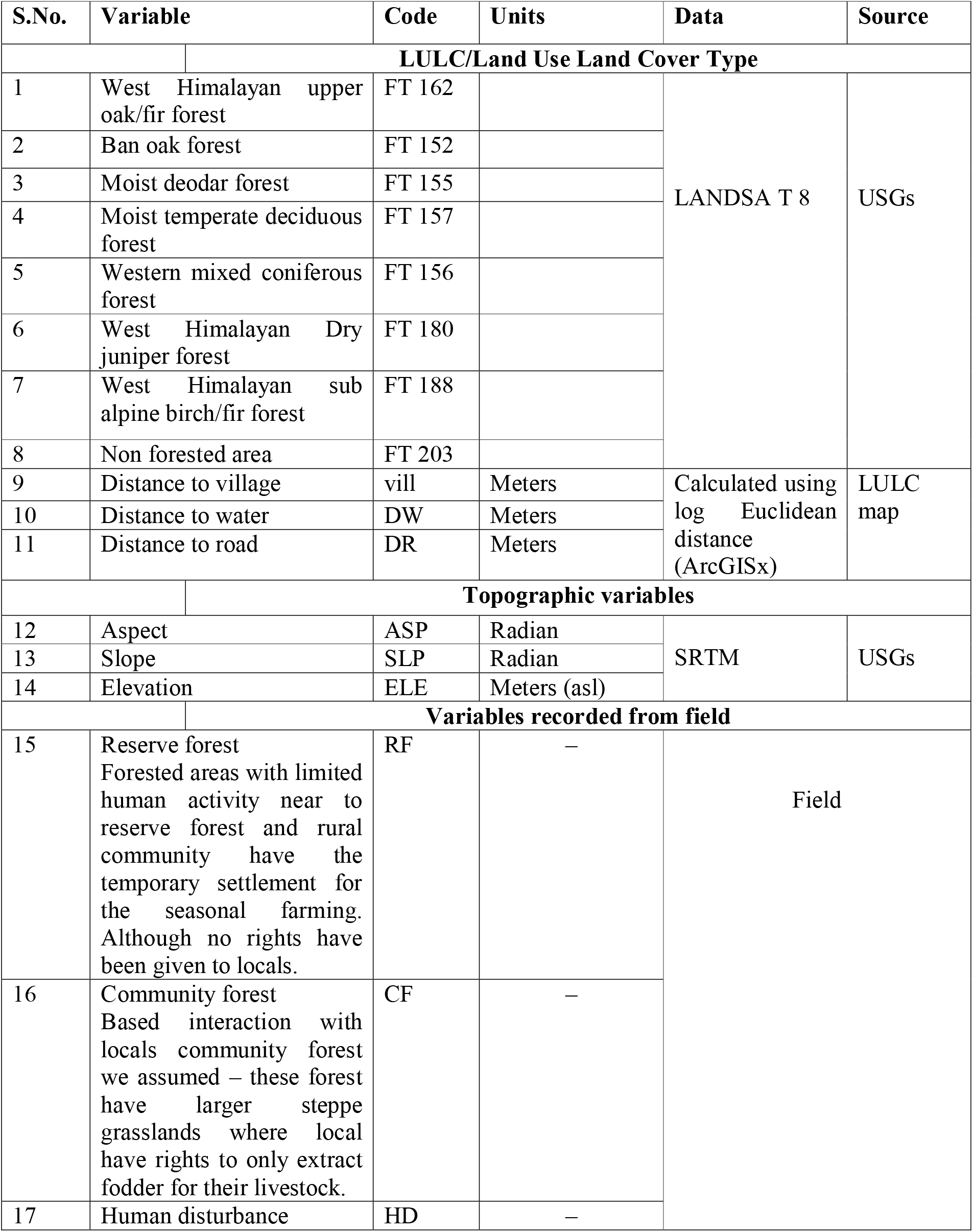
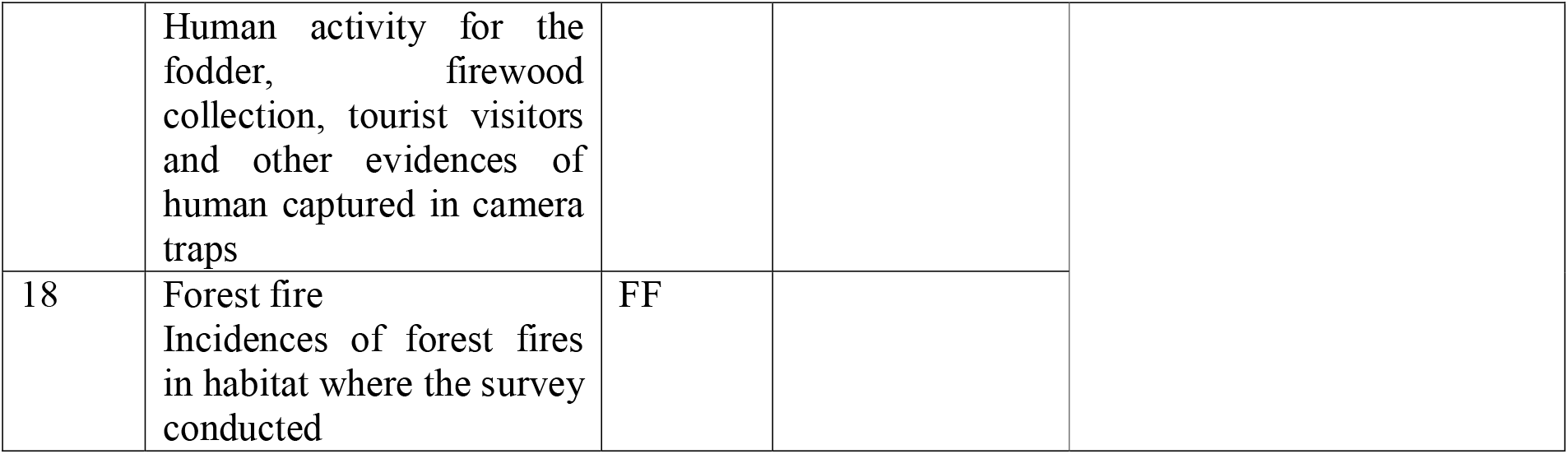
Habitat variables used for occupancy analysis of Himalayan monal in Uttarkashi, Uttarakhand.

| S.No. | Variable | Code | Units | Data | Source |
| --- | --- | --- | --- | --- | --- |
|  |  | LULC/Land Use Land Cover Type |  |  |  |
| 1 | West Himalayan upper oak/fir forest | FT 162 |  | LANDSAT 8 | USGs |
| 2 | Ban oak forest | FT 152 |  |  |  |
| 3 | Moist deodar forest | FT 155 |  |  |  |
| 4 | Moist temperate deciduous forest | FT 157 |  |  |  |
| 5 | Western mixed coniferous forest | FT 156 |  |  |  |
| 6 | West Himalayan Dry juniper forest | FT 180 |  |  |  |
| 7 | West Himalayan sub alpine birch/fir forest | FT 188 |  |  |  |
| 8 | Non forested area | FT 203 |  |  |  |
| 9 | Distance to village | vill | Meters | Calculated using log Euclidean distance (ArcGISx) | LULC map |
| 10 | Distance to water | DW | Meters |  |  |
| 11 | Distance to road | DR | Meters |  |  |
|  |  | Topographic variables |  |  |  |
| 12 | Aspect | ASP | Radian | SRTM | USGs |
| 13 | Slope | SLP | Radian |  |  |
| 14 | Elevation | ELE | Meters (asl) |  |  |
|  |  | Variables recorded from field |  |  |  |
| 15 | Reserve forest<br>Forested areas with limited human activity near to reserve forest and rural community have the temporary settlement for the seasonal farming. Although no rights have been given to locals. | RF | — | Field |  |
| 16 | Community forest<br>Based interaction with locals community forest we assumed – these forest have larger steppe grasslands where local have rights to only extract fodder for their livestock. | CF | — |  |  |
| 17 | Human disturbance | HD | — |  |  |

|  |  |  |
| --- | --- | --- |
|  | Human activity for the fodder, firewood collection, tourist visitors and other evidences of human captured in camera traps |  |
| 18 | Forest fire<br>Incidences of forest fires in habitat where the survey conducted | FF |

Furthermore, the selected grids were further divided into 5 km × 5 km grids, and out of the total 104 grids only 35 grids were surveyed. These selected 35 grids were further divided into 2 km × 2 km for the intensive sampling based on the movement ecology of the study species Li et al (2010); Selvan et al (2013). However, some of the grids within 2 km× 2 km were dropped because of the presence of sparse human settlements, non-forested areas and few of them were not logistically possible to cover (Fig. 2). These intensive sampling grids were systematically surveyed using 54 transects of varied length (1.5 to 7 km) of about total 650 km length representing all elevational gradient from 2200 m to 4067 m in the species distribution ranges and also placed 69 camera traps in these selected grids (Supplementary File 1; Figure S1).

**Fig. 2.**
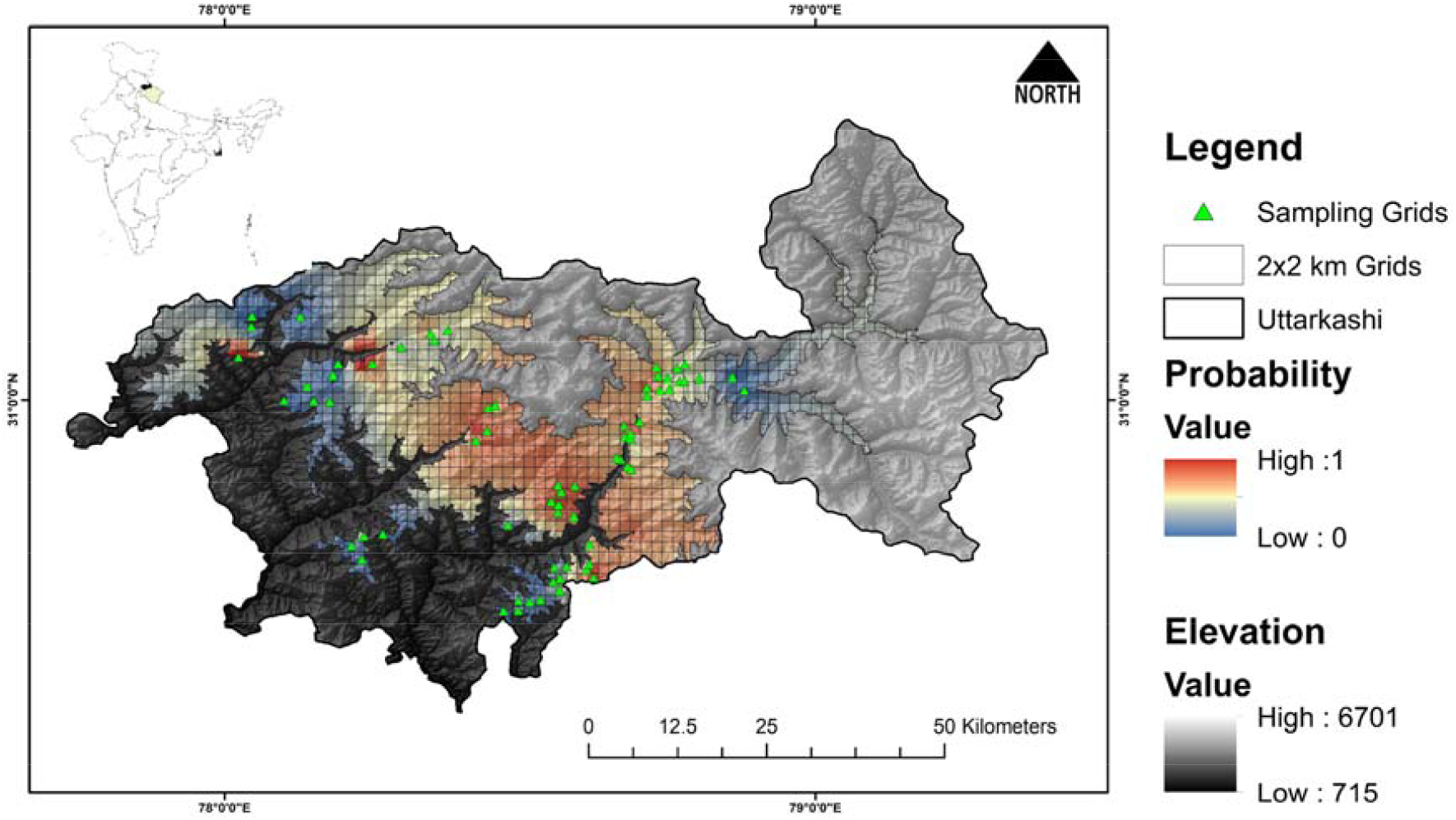
Sampling grids (2*2 km) for Himalayan monal in Uttarkashi.

For habitat characterization of the species, we laid 10 m radius plots around each camera locations and also at all sites were species sighted and an indirect sign were observed for vegetation sampling collection. Information such as GPS location, elevation, aspect, slope, habitat type, distance to the nearest water source, and distance to the village was recorded for all sites. The human disturbance was noted as the presence of locals in the area as to collect the fodder, firewood collection, and other activities on transects and the human captures were extracted from the camera traps. The camera traps were placed on animal trails and at about 1–2 feet height from the ground as optimized during reconnaissance survey based on the angle and coverage of species the montane ecosystem system. For capturing the presence, ultra-compact SPYPOINT FORCE-11D trail camera (SPYPOINT, GG Telecom, Canada, QC) and Browning Trail Camera (Defender 850, 20 MP, Prometheus Group, LLC Birmingham, Alabama, https://browningtrailcameras.com) were used. We divided our camera–trapping survey into seven temporal replicates (considering seven days as one occasion). The capture rates (total number of photographs captured/total number of camera trap days) are significantly correlated to abundance estimates of a species O’Brien et al (2003); Silveira et al (2003).

### Covariates

We hypothesized that habitat variables might influence the occupancy and detection probability of Himalayan monal in the study area. Hence, a total of 18 variables were extracted either from the field or using the ArcGIS v. 10.6 software (ESRI, Redlands, CA). These covariates were categorized as topographic, habitat type and anthropogenic variables (Table 1). The topographic variables (elevation, slope and aspect) were generated using 30 m resolution SRTM image downloaded from Earth Explorer (https://earthexplorer.usgs.gov/). The Landsat 8 satellite imagery (Spatial resolution = 30m) downloaded from Global Land Cover Facility was classified into seven habitat types following Singh et al (2005) using ERDAS Image version 9.0 (ERDAS®, Inc, Atlanta, Georgia) with classification accuracy 91.33%. The seven habitat types identified are West Himalayan Sub-alpine birch/fir Forest (FT 188), West Himalayan upper oak/fir forest (FT 162), West Himalayan Dry juniper forest (FT 180), Ban oak forest (FT 152), Moist Deodar forest (FT 155), Western mixed coniferous forest (FT 156), Moist temperate Deciduous forest (FT 157). All the variables were tested using the pairwise Spearman’s rank correlation coefficient (rs) and those significantly correlated were dropped from the further analysis (rs<0.7). These habitats were used for further analysis considering their importance to species ecology and behaviour. The values for all the covariates were extracted at 30 m resolution. A single value per site was obtained by averaging all the pixel values within each sampling site.

### Occupancy framework

Camera trap data and sign survey data was used as an index of habitat use by the Himalayan monal based on their occupancy status in a given sampling grid. All continuous variables were z-standardized before proceeding to analysis. Z-transformation accounts continuous variables and compared the scores from disparate distribution in which the mean of the population will be 0 and the standard deviation 1 which allow the comparison of dissimilar metrics Hines (2006). Since, the detection of Himalayan monal was likely to be imperfect an occupancy framework was used to address this issue MacKenzie et al (2002); Mackenzie & Royle (2005); Mackenzie et al (2006). In the present study, we adopted single-season occupancy analysis for estimating the site occupancy probability (ψ) and detection probability (ρ) of Himalayan monal using a likelihood-based method described by MacKenzie et al (2002). The analysis was carried out using the PRESENCE v.2.12.25 software package Hines (2006). The detection/non-detection histories for each grid were prepared from camera trap surveys as well as from sign survey over a study period of 12 months. The surveys were divided into seven sampling occasions of 7 days each. We pooled the data from all the locations at the respective sites and constructed standard detection histories for each site Mackenzie et al (2006).

A logit link function Mackenzie et al (2006) was used to model Himalayan monal presence as dependent on habitat covariates in program PRESENCE v.2.12.25 Hines (2006). The two-stage logistic regression analysis was performed to determine the effect of different site covariates on the Himalayan monal occupancy. We selected the habitat variables ecologically important for Himalayan monal and therefore, a number of models were developed, including each of the variables modelled separately as well as a combination. A total of 38 models were developed in which we used detection probability as a constant p(.) and allowed occupancy to vary ψ(covariate) with site-specific covariates. In the second step, we allowed occupancy ψ(.) as constant and detection probability p(covariate) as function to vary with the site-specific covariates MacKenzie et al (2006); Supplementary file 1; Table S1). All the candidate models were ranked based on their AIC (Akaike Information Criterion) values, and the Akaike weights Burnham & Anderson (2002). The Akaike weight represents the ratio of ∆AIC values for the whole set of candidate models, providing a strength of evidence for each model. Based on the AIC weight values, only the top-ranking candidate model was used to determine the Himalayan monal occupancy in the study area. The summed model weight of each covariate in these models was used to determine the most influential variables for the species. The sign of the logistic coefficient of each variable (positive or negative) was used to determine the direction of influence of the variables on the occupancy and detection probability of Himalayan monal in Uttarkashi. Further for interpolation in the entire range in the Uttarkashi district, a continuous occupancy probability surface was generated based on average site occupancy estimated of the species and by using Inverse Distance Weighted (IDW) interpolation technique in ArcGIS 10.6 Liszka (1984); Curveira-Santos et al (2019). Thus, it resulted in a spatial interpolation surface occupancy over the habitable elevation range of the species, i.e. between 2000m – 4200m elevation gradient.

### Abundance and density estimation

The abundance estimation approaches developed for occupancy surveys incorporate detection probability directly into the estimation process MacKenzie et al (2002). However, density estimation is difficult for species such as Himalayan monal for which individuals cannot be identified because of the lack of a distinct marking Singh et al (2014). In such cases, abundance estimates can be made with occupancy surveys that rely on a species being detected, or not, at a particular site MacKenzie et al (2002). We assumed that detection of individuals was independent, individuals were equally detectable across the whole sampling site, and the site-specific abundance of individuals followed a Poisson distribution Mackenzie et al (2006); Singh et al (2015). The Royle–Nichols model provides estimates of the abundance (λ) and detectability (r), representing the average abundance per site and innate species detectability, respectively Royle & Nichols (2003). We adopted this approach and divided abundance (λ) by the area of the sampling unit (n=69; area=276 km^2^) to estimate the average Himalayan monal density of the sites in the study area. We used the Royle–Nichols heterogeneity constant model λ(.),r(.) to estimate the abundance and associated parameters of Himalayan monal following Singh et al (2015)

### Activity pattern

The daily activity pattern of Himalayan monal and the overlap between the temporal patterns of Himalayan monal and humans sharing the same habitat were also estimated. We have compared activity pattern of Himalayan monal and humans to understand the overlapping patterns using package “Overlap” in R 3.5.1 (R Development Core Team) and assess how the human activity is influencing the daily activity pattern of Himalayan monal. The time and date printed on the photographs have been used to determine the daily activity pattern of individual species Pei (1998). We used a Daily Activity Index (DAI) of half an hour duration to examine the daily activity (Dinata et al. 2008). The coefficient of overlap is denoted by “Dhat1” values, ranging between zero (no overlap) and 1.0 (complete overlap).

## Result

The total effort of 2819 camera trap nights in 69 camera traps and 54 trails covering 650 km of different locations resulted in 40 independent captures of Himalayan monal (*Lophophorus impegans*) through camera trapping and 99 independent signs recorded through sign survey in the study area. The Himalayan monal occupies the elevational range between 2000 to 4067 m in the study area and mostly concentrating in a narrow belt of 2400 – 3400 m (Fig. 3d).

**Fig. 3.**
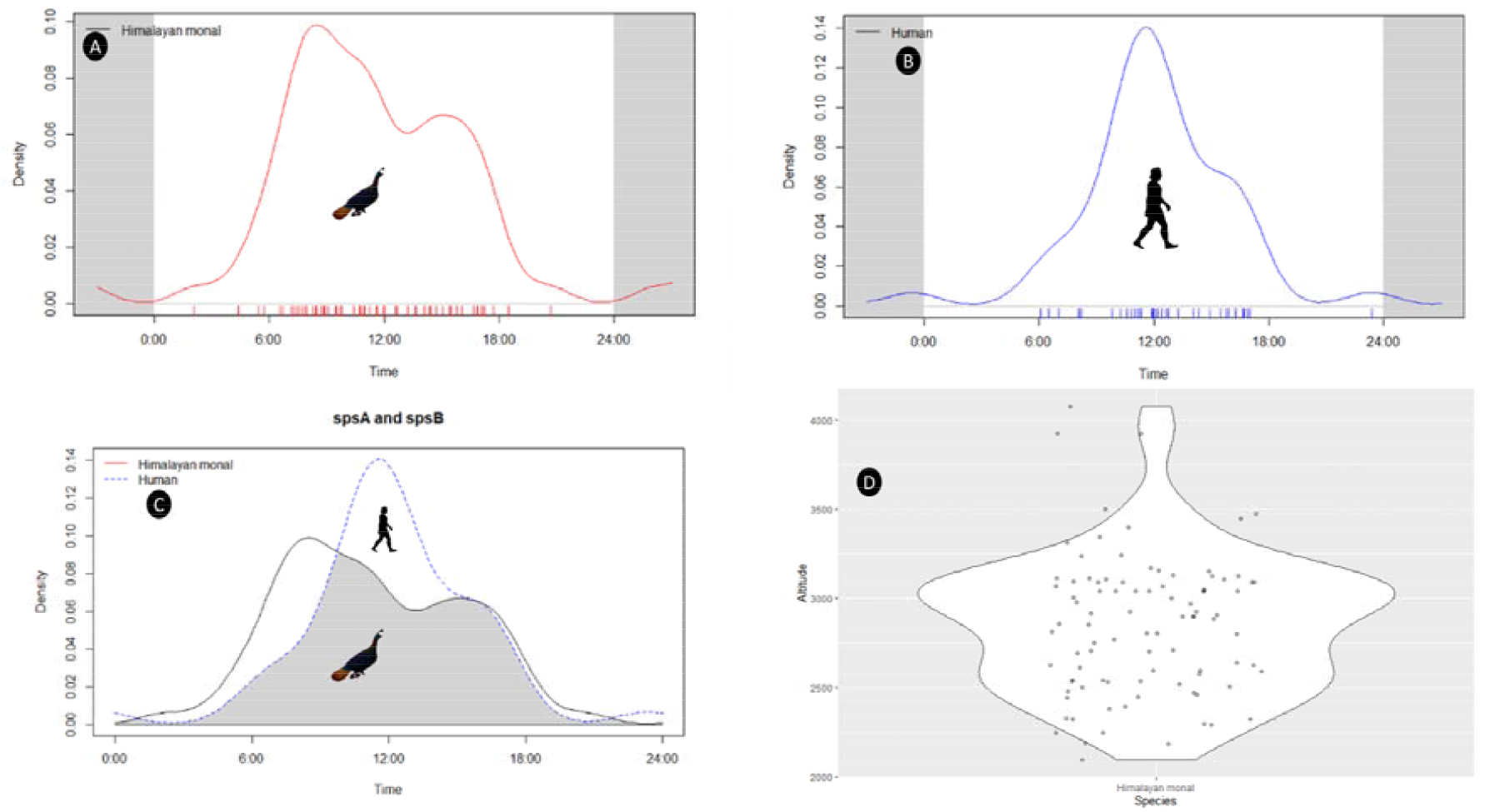
Temporal activity pattern of Himalayan monal (a), human (b) and overlapping activity pattern of Himalayan monal (c), and violin plot depicts elevational distribution of Himalayan monal (d) in Uttarkashi.

The combination of both, i.e., camera traps and sign survey yielded with 0.69 naïve occupancy and the mean estimated occupancy of monal ranged from 0.1 to 1.0 (Table 2, Fig. 1). Out of 38 models, only five top models are shown, among which top model were found useful in explaining the influence of covariates on monal occupancy and detection probability based on the lowest AIC values (Table 2). The top model based on the AIC value indicated that the slope and reserve forest (RF) influenced the occupancy of Himalayan monal the most (Table 3) while keeping detection probability constant p(.). This model assumed the occupancy of the Himalayan monal at different sites varied as a function of slope and reserve forest. The results of our top model suggest that occupancy probability of monal were positively related to landscape slope (β =27.52± SE 16.25) and negatively influenced by reserve forest (RF) (β= −8.14±SE 4.99) (Table 4).

**Table 2:** Occupancy estimates of Himalayan monal in different forest ranges of Uttarkashi, Dehradun.

| **Codes** | **$\Psi$** | **±SE** | **95% CI** |
| --- | --- | --- | --- |
| Gangotri | 0.98 | 0.02 | 0.007-0.07 |
| Taknor | 1 | 0.00 | 1-1 |
| Badahat | 0.99 | 0.001 | 0.75-1 |
| Mukhem | 0.77 | 0.06 | 0.51-0.88 |
| Darasu | 0.79 | 0.09 | 0.46-0.94 |
| Yamnotri | 1 | 0 | 1-1 |
| Singtur | 0.11 | 0.085 | 0.02-0.42 |
| Rupin | 0.67 | 0.06 | 0.02-0.77 |
| Sankari | 0.84 | 0.03 | 0.65-0.91 |

$\Psi$ – Occupancy probability; ± SE – Standard error; CI – Confidence intervals

**Table 3:** Summary of top five models selected for Himalayan monal occupancy in Uttarkashi, Uttarakhand Ψ – occupancy probability; p – detection probability; Codes are explained in Table 1; K is number of parameters

| Model | AIC | Delta AIC | AIC weights | Model likelihood | K |
| --- | --- | --- | --- | --- | --- |
| $\Psi(\text{Slope+RF}), p(.)$ | 391.91 | 0 | 0.58 | 1 | 3 |
| $\Psi(\text{Slope}), p(.)$ | 393.76 | 1.85 | 0.23 | 0.39 | 2 |
| $\Psi(\text{RF}), p(.)$ | 396 | 4.09 | 0.076 | 0.12 | 2 |
| $\Psi(\text{Slope+RF}), p(\text{RF})$ | 398.1 | 6.19 | 0.02 | 0.04 | 3 |
| $\Psi(.), p(\text{RF})$ | 400.66 | 8.75 | 0.007 | 0.01 | 2 |

**Table 4:** Models with beta coefficient values of different site covariates influencing Himalayan monal occupancy and detection probability. Ψ – occupancy probability; p – detection probability; Codes as explained in Table 1.

| **Model** | **$\psi$ (Slope)** | **$\psi$ (RF)** | **$\Psi$(.)** | **p (.)** | **p(RF)** |
| --- | --- | --- | --- | --- | --- |
| psi(Slope+RF),p(.) | 27.52±16.25 | -8.14±4.99 | - | -1.29±0.13 | - |
| psi(Slope),p(.) | 3.60±1.30 | - | - | -1.20±0.16 | - |
| psi(RF),p(.) | 1.74±0.65 | - | - | -1.18±0.67 | - |
| psi(Slope+RF),p(RF) | 1.08±1.37 | 1.22±1.00 | - | - | 1.22±0.18 |
| Psi(.),p(RF) | - | - | - | 1.25±0.38 | -1.16±0.16 |

### Abundance and density estimation

The estimated site mean occupancy of Himalayan monal in the Uttarkahsi was 0.82±0.08 with a detection probability of 0.23±0.03 (i.e., probability of detection of Himalayan monal on each survey) using the null model ψ (.) ρ (.) in which occupancy and detection probability are kept constant. Our results showed that the estimated occupancy of Himalayan monal was higher than naïve occupancy (ψ naive=0.69; ψ estimated=0.82) (Table 5). The abundance of Himalayan monal in Uttarkashi based on Royle-Nicholas heterogeneity null model λ(.), r(.) was λ=2.48±1.48 monal/site with a detection probability of r=0.07±0.04. The overall abundance was 171.58±10.2 monals, with density estimated to be 0.62/km^2^ (171.58/276=0.62 km^2^) in the study area (Table 5).

**Table 5:** Occupancy and abundance estimation of Himalayan monal in Uttarkashi. (Density calculated in 2×2 Km grids).

| Parameters | Estimation |
| --- | --- |
| Sampled area | 276 km<sup>2</sup> |
| Overall abundance | 171.58±10.2 |
| Naïve site occupancy | 0.69 |
| Site occupancy predicted | 0.82±0.08 |
| Detection probability of Himalayan monal | 0.23±0.03 |
| Density | 0.62/ km<sup>2</sup> |

### Activity pattern

The daily activity pattern of Himalayan monal and Humans on visualization showed a marked difference in the peaks of their activity (Fig. 3). The Himalayan monal showed higher activities in the hours after sunrise, and it gradually decreased towards mid-day and gave a smaller peak before sunset and remains a little active till 18:00 hrs (Fig. 3). Whereas, the human activity in the forest was mostly restricted to hours with daylight, i.e., from 05:47 to 18:26 hrs and showed greater peak activity in mid-day. The overlap plot of temporal activity between Himalayan monal and humans showed a marked difference in the activity of Himalayan monal due to the presence of humans (Overlap coefficient–Dhat1=0.77; Fig. 3).

## Discussion

The distribution and habitat use assessment information is often required for making informed conservation and management decisions. The Himalayan monal is one of the least studied species in India, and whatever is known is mainly available in the form of short term surveys and status reports(Ramesh et al (1999); Bhattacharya et al (2009); Miller (2010); Jolli & M. K. Pandit (2011); WII (2016). However, the majority of the studies were restricted to the PAs only, and no information is available from habitats of species outside the PA network. Here, for the first time, we attempted to understand the occupancy and abundance of monal in Uttarkashi primarily outside the PA network. We have modelled the occupancy of monal by explicitly incorporating detection probability and habitat variables assuming that these variables may have an ecological impact on species distribution and occupancy.

The Himalayan monal although is listed among the Least Concern species IUCN (2020), but has been provided maximum protection as Schedule I of WPA 1972 considering the threats species is facing because of illegal poaching and habitat loss. The populations of this spectacular bird is declining throughout its distribution range because of different anthropogenic threats (Ramesh (2003); Chen et al (2018). The species is hunted in many areas because of its plumage and meat Fuller & Garson (2000); Selvan et al (2013). Although the species is losing ground in many areas not much effort has been made towards developing knowledge on its various ecological aspects, however, what is known today is precisely a decade old and restricted to few PAs Ramesh et al (1999b); Great Himalayan National Park, Himachal Pradesh; Hussain et al (2001) (Hussain et al. 2001) in Kumaon Himalayas, Uttarakhand; Hussain & Sultana (2013) in Kumaon Hills, Uttarakhand; Jolli & Pandit (2011a) Western Himalaya. Especially no study has been conducting in non-protected forests of Uttarkashi, even though they are present in good numbers despite these forest ranges are anthropogenically disturbed. Based on the top model results indicated that the occupancy of Himalayan monal is best predicted by the slope (β =27.52±16.25) and reserve forest (β= −8.14±4.99) (Table 4). It is quite plausible that Himalayan monal is preferring steeper slopes for roosting as a strategy to escape depredation by carnivores and also steeper slopes are devoid of anthropogenic disturbances. Other studies Beebe (1922); Hussain & Sultana (2013) also corroborated with our findings which highlighted that habitat preference of the species is positively influence by the slope.

Further, the negative association of reserve forest with the occupancy of the species indicates that Himalayan monal avoids human-disturbed areas, as the species is susceptible to human disturbance. Our results further strengthen the findings of studies indicating that the species is sensitive to anthropogenic disturbances and therefore species might have developed a behavioural tendency of avoiding areas with anthropogenic disturbances Ramesh (2003); Bhattacharya et al (2009); Jolli & M. K. Pandit (2011). Moreover, the overlapping activity pattern of Himalayan monal and humans also indicates that Himalayan monal restricts its daily activity when humans are most active (Fig. 3).

The naive occupancy of 0.69 indicates that the forest ranges in the study landscape have the potential for the conservation of Himalayan monal irrespective of the fact that the majority of the forests in the study areas are outside the PAs. However, the Himalayan monal was not always detected when they were present at a site indicated by lower detection probabilities (p=0.23±0.03). The density of Himalayan monal in the study area was estimated to be 0.62 individuals/km^2^ which is relatively lesser than the reported densities from Great Himalayan National Park (2.87 ± 0.22 birds/km, to 8.0 birds/km: Gaston & Garson (1981); Jolli & Pandit (2011b); Miller (2010) and the buffer zone of Nanda Devi Biosphere reserve (36.37 ± 2.69 individuals/km^2^: Bhattacharya et al (2009) and this difference may be due to different methods and areas. The higher densities reported earlier were from PAs which are better protected and managed with the aim of in-situ conservation of wildlife species. Hence, our results cannot be compared with them as the present study landscape is not managed under the protection regime instead focus of the management is production forestry and the forest stands are managed under various silviculture working plans. Furthermore, earlier studies were carried out in best-known habitats of the species, and the estimates were largely based on trails observations. In Uttarkashi, the abundance was estimated to be λ=2.48±1.48 monal/site with the overall abundance of 171.58±10.2 Himalayan monal, in the entire study areas.

The violin plot depicts that the Himalayan monal occupies the elevation range between 2000 to 4067 m in the study area (Fig 3d). It occupies upper temperate oak-conifer forests, subalpine oak forests interspersed with open grassy slopes, cliffs and alpine meadows, but mostly concentrating in a narrow belt of 2700 – 3700 m Grimmett et al (1998). However, in the present study majority of the captures and indirect observations were from 2400-3400 m. In winter Himalayan monal moved to lower altitudes, and they exhibit apparent altitudinal migration reaching as low as 2000 m in winter Ramesh (2003).

Although the species has an extensive range in the Himalayas, it is getting vulnerable to poaching, habitat loss and degradation Hoyo et al (1994); Miller (2010). This conservation priority bird is facing declines in population primarily due to fragmentation and degradation of the habitat, and poaching for local consumption, especially in winter when the birds descend close to human habitation Ramesh (2003); Poudyal et al (2013). Hence, maintaining insurance populations of this charismatic species thus becomes imperative. The results of our study suggest that Himalayan monal is extremely sensitive to human disturbances and avoids human disturbance areas. Further, Jolli & Pandit (2011a) highlighted that increasing human interference is negatively impacting the populations of pheasants in the Himalayan region.

## Conclusion

The systematic information on the species with high conservation values required a better understanding of their ecology, behaviour and distribution. The present study provides information on distribution, occupancy, activity pattern and abundance of monal primarily from non-protected forest ranges of Uttarkashi district, which was not known earlier and can be replicated for the other species in the similar ecosystem. We found that habitat sampled outside the PA network holds a good population of species which is good for its long-term viability. The sites with higher occupancy probability of Himalayan monal in Uttarkashi (Fig 4) should be prioritized for long term monitoring. As these sites can be effective for expressing breeding behaviour associated with their microhabitat requirement Kajin Eduardo & Grelle (2012). These sites may also be identified for enhancing protection in addition to the PAs by adopting landscape conservation strategy. Hence, we recommend landscape-level conservation actions for securing the future of the species, as landscape approach will enhance gene flow among the populations and also mitigate the impacts of habitat fragmentation. Moreover, the information generated in the present analysis sets a benchmark which will be imperative for the future monitoring of the species population.

## Acknowledgement

We thank Chief Wildlife Warden, Forest Department Uttarakhand, and Government of Himachal Pradesh for granting the necessary permission to undertake field surveys the research work. Authors also thanks to the Director, Head of Office and O/C Technical Section, Zoological Survey of India for consistent support during the study. Authors are thankful to Shree Sandeep Kumar, Divisional Forest Officer (DFO), Uttarkashi Forest Division for his consistent guidance and support during the fieldwork. Authors are extending thanks to DFOs of Tons Forest Division, Purola and Upper Yamuna Forest Division, Badkot, Deputy Director, Govind Pashu Vihar National Park and Sanctuary, Purola for helping and guiding the field team of ZSI, Kolkata time to time. We are thankful to all the Range Officers and field staff of all three Forest Divisions, National park for their active support and logistics during the fieldwork. We acknowledge the National Mission for Himalayan Studies, Ministry of Environment, Forest and Climate Change (MoEF&CC) for the funding support under the Grant No. NMHS/2017-18/LG09/02.

